# Large-scale study validates that regional fungicide applications are major determinants of resistance evolution in the wheat pathogen *Zymoseptoria tritici* in France

**DOI:** 10.1101/2020.07.17.208728

**Authors:** Maxime Garnault, Clémentine Duplaix, Pierre Leroux, Gilles Couleaud, Olivier David, Anne-Sophie Walker, Florence Carpentier

## Abstract

- Research rationale: In modern cropping systems, the near-universal use of plant protection products selects for resistance in pest populations. The emergence and evolution of this adaptive trait threaten treatment efficacy. We identified determinants of fungicide resistance evolution and quantified their effects at a large spatiotemporal scale.
- Methods: We focused on *Zymoseptoria tritici*, which causes leaf blotch in wheat. Phenotypes of qualitative or quantitative resistance to various fungicides were monitored annually, from 2004 to 2017, at about 70 sites throughout 22 regions of France (territorial units of 25 000km^2^ on average). We modelled changes in resistance frequency with regional anti-*Septoria* fungicide use, yield losses due to the disease and the regional area under organic wheat.
- Key results: The major driver of resistance dynamics was fungicide use at the regional scale. We estimated its effect on the increase in resistance and relative apparent fitness of each resistance phenotype. The predictions of the model replicated the spatiotemporal patterns of resistance observed in field populations (R^2^ from 0.56 to 0.82).
- Main conclusion: The evolution of fungicide resistance is mainly determined at the regional scale. This study therefore showed that collective management at the regional scale could effectively complete local actions.

## 1 INTRODUCTION

The efficacy of pesticides and drugs has been compromised by the rapid and widespread evolution of resistance, increasing their use to maintain control (Georghiou & Mellon, 1983; Russell, 2005; Gould *et al*., 2018). Resistance management is essential for human health, biodiversity and food security, given the rapid emergence and spread of resistance and the lack of new modes of action (MoA) (Palumbi, 2001; Grimmer *et al*., 2014). Many studies have investigated the effects of various factors on the evolution of resistance: fitness cost (Andersson, 2003), mutation rate (Martinez & Baquero, 2000; Gressel, 2011), population size (Sisterson *et al*., 2004), strength of selection pressure and its mitigation in strategies (Oz *et al*., 2014; van den Bosch *et al*., 2014). A number of studies have advocated further work on the relative impact of these factors on a given pest and of the interactions between these factors (Berendonk *et al*., 2015; Hughes & Andersson, 2015), with a view to promoting large-scale strategies (Okeke *et al*., 2005; Menalled *et al*., 2016).

Several studies have shown how agricultural selection pressures affect the large-scale structure of pest populations at national scale. For instance, the national distribution of resistance varieties shapes the adaptation of pathogen populations to cultivars (Tyutyunov *et al*., 2008; Papaïx *et al*., 2011). Historical herbicide applications drive the evolution of herbicide resistance at a national scale (Hicks *et al*., 2018). For fungicide resistance, theoretical studies have revealed that combining effective MoAs over time and space can delay resistance evolution (REX Consortium, 2013; van den Bosch *et al*., 2014) and that large-scale management strategies may differ from and interact with in-field strategies (Parnell *et al*., 2006). However, so far, in the absence of large-scale studies, recommendations about fungicide use mostly stem from empirical studies conducted in local field trials assessing the impact of different spraying strategies (Rosenzweig *et al*., 2008; Dooley *et al*., 2016a,b; Heick *et al*., 2017). In France, the metropolitan area was until 2016 subdivided into 22 administrative regions. Their average area of 25,000 square kilometres is consistent with the long-distance dispersal capabilities of pathogens, e.g. between ten to hundred kilometres for airborne fungi (Brown & Hovmøller, 2002; Linde *et al*., 2002). Homogeneous agronomic properties and pedoclimatic conditions generally lead to similar crop sequences in regions (Xiao *et al*., 2014). Therefore, this scale seems relevant for both studying the evolution of resistance and implementing large-scaled resistance management. The aim of this study was to highlight the determinants at the regional scale of fungicide resistance evolution in France. First, we considered (i) the regional use of fungicides, since the quantity of fungicide sprayed increases the selection of resistances (van den Bosch *et al*., 2014), (ii) the refuge effect (Parnell *et al*., 2006; Tabashnik *et al*., 2008) by including the area under organic farming, as a proxy of unsprayed fields, since the heterogeneity of the selection may reduce the resistance growth (REX Consortium, 2010, 2013) (iii) the population size, using yield losses as a proxy, because it may affect the evolution of resistance alleles through genetic drift (Maxwell *et al*., 1990; Sisterson *et al*., 2004). We did not consider local field-scaled strategies (e.g. adjustment of spray timing, fungicides mixture or alternation).

We focused our analysis on *Zymoseptoria tritici* (formerly *Septoria tritici* and *Mycosphaerella graminicola* as teleomorph), an ascomycete responsible for septoria leaf blotch (STB) on winter wheat. *Z. tritici* has many features facilitating the emergence of resistance: high genome plasticity, a large population size, high genetic diversity, asexual and sexual reproduction, an ability to disperse over large distances (Zhan & McDonald, 2004; Croll & McDonald, 2012). STB is a major wheat disease that can cause yield losses of up to 50% (Ponomarenko *et al*., 2011; Torriani *et al*., 2015). In western Europe, up to 70% of all fungicide use is linked to STB control (Fones & Gurr, 2015). As a result, various degrees of resistance to all authorised unisite inhibitors (*i*.*e*. exhibiting a single molecular mode of action) have been observed in France (Garnault *et al*., 2019).

We previously published an initial analysis of the Performance trial network dataset, in which phenotypes of resistance to four fungicide MoAs were monitored annually, from 2004 to 2017, at about 70 sites throughout France (Garnault *et al*. 2019). We found significant differences between phenotypes in terms of changes in spatial distribution and/or growth rates. Major differences in population structure and dynamics were highlighted between the North and South of France.

We develop here an explanatory model for identifying the determinants of these regional spatiotemporal heterogeneities in resistance evolution according to resistance phenotype. We investigated the effect of three potential drivers of selection pressure through the (*i*) regional fungicide uses and (*ii*) fraction of sprayed fields, and we also considered (*iii*) the effect of the pathogen population size which may affect evolution through genetic drift. Our analysis shows that the inter-annual change in resistance frequency can be assessed at regional scale, and that the major determinant of resistance is the selection pressure exerted by fungicide applications in the preceding year. This study provides empirical results for regional resistance management, at a level intermediate between field and national recommendations. A sound understanding of resistance evolution and of its determinants would help optimizing resistance management and applying them at sound spatiotemporal scales. It should ultimately help to reduce pesticide use in agrosystems.

## 2 MATERIALS AND METHODS

### 2.1 Data description

#### 2.1.1 Sampling of *Z. tritici* populations and estimation of resistance frequency

The “Performance network” is supervised by ARVALIS-Institut du Végétal and the INRAE research institute at Thiverval-Grignon. It carried out field trials on wheat throughout France between 2004 and 2017 with a mean of 70 trials annually (4 to 5 trials per region and year on average; 90% credible interval is 1 to 10). The frequency of resistant phenotypes in *Z. tritici* populations sampled annually in these trials is recorded in the associated dataset (see Garnault *et al*., 2019 for further information).

Wheat trials were carried out in a randomized block design with 3 to 4 replicates. The frequency of resistant phenotypes in populations was estimated by collecting bulk pycnidiospores from 30 to 40 upper leaves that were randomly sampled within each plot and showed STB symptoms. Cropped cultivars were predominantly STB-sensitive to promote the presence of the disease. A total of 124 different wheat cultivars were cropped over the dataset. Phenotypes were distinguished on the basis of their germination or growth on Petri dishes containing discriminatory doses of fungicides, optimised on individual genotyped isolates (see Leroux & Walker, 2011 and Garnault *et al*., 2019 for more details). We then considered: (*i*) the phenotype displaying specific qualitative resistance to strobilurins (or QoIs; inhibitors of respiration complex III), hereafter referred to as the StrR phenotype, (*ii*) the group of phenotypes with moderate quantitative resistance to DMIs (sterol 14α-demethylation inhibitors), hereafter referred to as TriMR phenotypes, (*iii*) the group of phenotypes with a high quantitative resistance to DMIs, hereafter referred to as TriHR phenotypes. The TriMR group encompasses the TriR6 and TriR7-TriR8 phenotypes, which were also included in the analysis (TriR6 strains grew on low doses of prochloraz, contrasting with the lack of growth of TriR7–TriR8 strains in these conditions; Leroux & Walker, 2011).

Region, year, sampling date and cultivar grown were recorded for each sample. We considered only populations from unsprayed plots for this study. The plots were sampled two times: at “S1” in April-May, at about the Z32 wheat stage (*n*=1320, from 2006 to 2011), and at “S2” in May-June, at about the Z39-Z55 wheat stage (*n*=2407, from 2004 to 2017).

In this study, we focused on resistance selection (*i*.*e*. after emergence and before complete fixation). For each phenotype, we defined a time interval grouping the years for which at least a quarter of the regions exhibited frequencies differing from 0% and 100%. We excluded the years following the “decline” of resistance, identified as the year when the frequency of resistance was no longer significantly higher than the average frequency. These frequency differences were estimated in Garnault *et al*. (2019). The year 2012 was the first year of decline for TriMR (other phenotypes did not show a decline). Time periods were respectively 2004 to 2012 (*n*=852, 16 regions) for the StrR phenotype, 2005 to 2011 (*n*=754, 16 regions) for the TriMR phenotype group, and 2010 to 2017 (*n*=360, 14 regions) for the TriHR phenotype group. We also included data from 2006 to 2017 for the TriR6 and TriR7-TriR8 phenotypes (*n*=910 and *n*=851, respectively), for analysis of the spatial heterogeneity of their frequencies. The data are summarised in Table 1.

**Table 1.** Data from the Performance network database used for the statistical analysis. Each observation corresponds to a resistance frequency measurement. “First year observed”: first year of observation in the Performance network. “Number of years observed”: total number of successive years of monitoring, including the first year. “Number of regions monitored”: total number of sampled French regions. “Number of regions monitored yearly”: total number of French regions with at least one observation each year. All regions were monitored for at least 80% of the studied time range. The last column sums up the total number of observed frequencies for each phenotype.

| Phenotype | First year<br>observed | Number of years<br>observed | Number of regions<br>monitored | Number of regions<br>monitored yearly | Number of<br>observations |
| --- | --- | --- | --- | --- | --- |
| StrR | 2004 | 9 | 16 | 9 | 852 |
| TriMR Total | 2005 | 7 | 16 | 13 | 754 |
| TriR6 | 2006 | 12 | 16 | 6 | 910 |
| TriR7-TriR8 | 2006 | 12 | 16 | 6 | 851 |
| TriHR | 2010 | 8 | 14 | 6 | 360 |
| Total |  |  |  |  | **3727** |

#### 2.1.2 Regional fungicide use

Every year, Bayer Crop Science uses field surveys to estimate the area of wheat sprayed with fungicides containing anti-STB active ingredients (AIs) in each region. These data do not include dose information but provide information about the areas sprayed with each AI and the number of sprayings (areas result from the multiplication of these two values). These areas are expressed in deployed hectares.

We retained the most widely used AIs for each MoA (AIs accounting cumulatively for more than 95% of the use of the MoA), to prevent background noise from AIs with a limited impact on STB control. The model therefore included pyraclostrobin (26%), azoxystrobin (19%), trifloxystrobin (15%), kresoxim-methyl (15%), fluoxastrobin (12%) and picoxystrobin (12%) for QoIs; and epoxiconazole (30%), prochloraz (17%), tebuconazole (13%), cyproconazole (11%), prothioconazole (10%), propiconazole (7%), metconazole (7%), fluquinconazole (2%) and hexaconazole (1%) for DMIs.

We took the regional heterogeneity in wheat production between regions (and, hence, in the area sprayed with fungicides) into account, by dividing the number of deployed hectares by the regional area under conventionally farmed wheat. The latter was calculated by subtracting the area under organic wheat (see section 2.1.4) from the total area under wheat (from the AGRESTE online data: agreste.agriculture.gouv.fr) for each year and region. This new variable unit was named 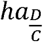 (*D* for deployed and *C* for cropped hectares), and was proportional to the mean number of times each AI was used over a cropping season in a given region. The national trend and the regional heterogeneity of fungicide use expressed in 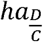 are shown for DMIs and QoIs in Fig. 1. Henceforth, this variable is denoted *F*_*itf*_, with *f* corresponding to the AI, *t* to the year and *i* to the region.

**Fig. 1.**
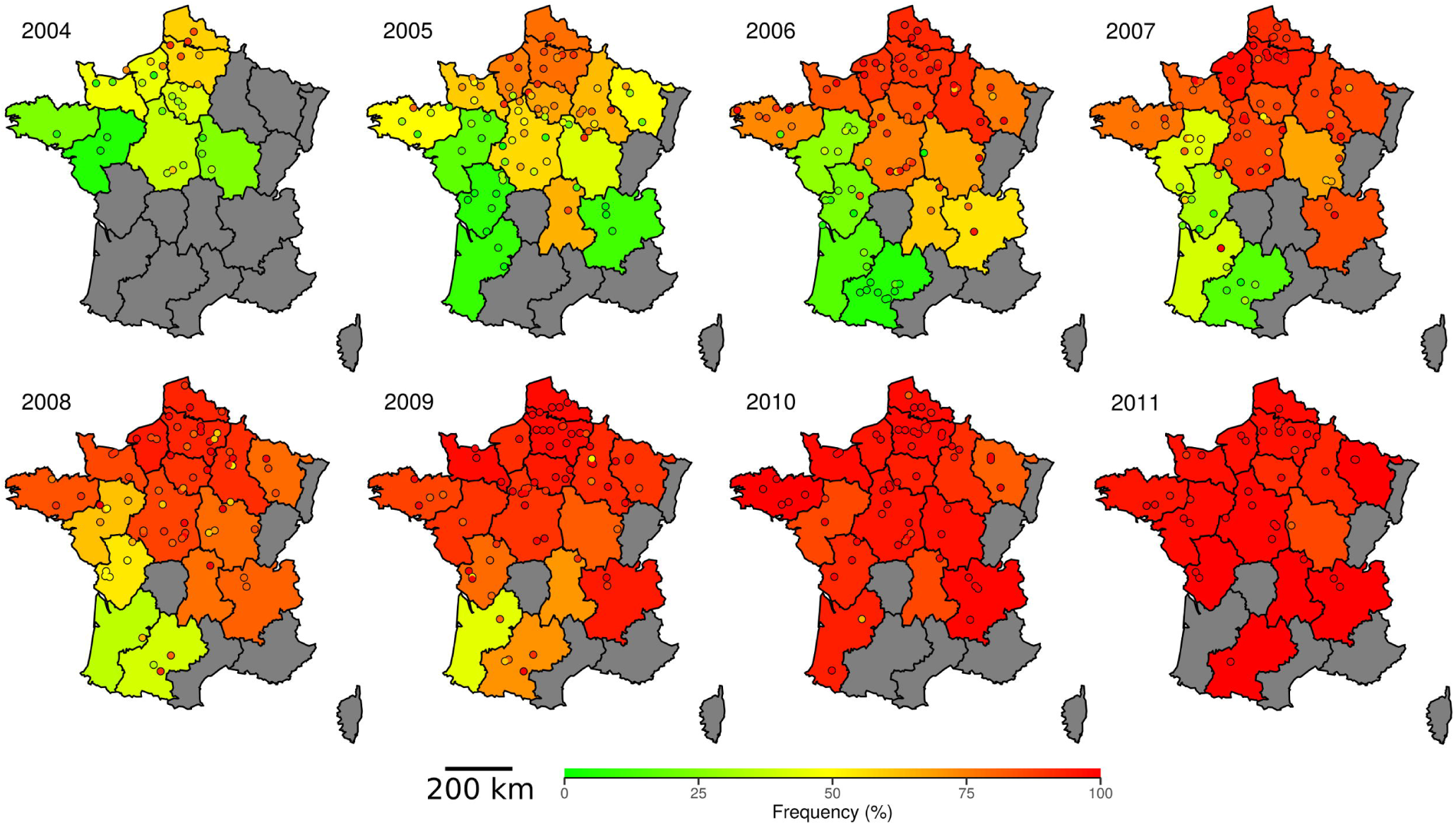
Changes in resistance frequencies (left) and explanatory variables (right, top to bottom, fungicide use, yield losses and area of wheat under organic farming). Bold lines: mean regional values. Shaded areas: quantiles of regional values (*i*.*e*. regional variability), 25% and 75% (dark grey), 2.5% and 97.5% (light grey). Dotted lines: regional minimum and maximum values. Fungicide use is expressed in 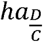 corresponding to the mean number of times each mode of action was used for spraying over a cropping season, regardless of the dose used. Yield losses are expressed in decitons (quintals) per hectare.

#### 2.1.3 Yield losses induced by STB

ARVALIS-Institut du Végétal assessed annual yield loss by conducting paired plot experiments throughout France with a mean of 80 trials (21 observations per region and year on average, distributed among 3 to 4 trials; 90% credible interval is 9 to 48), from 2004 to 2017, in 20 French regions (Arvalis, 2019). In each trial, we considered modalities cropped with STB-susceptible wheat cultivars, in both unsprayed plots and sprayed plots (providing maximum protection against diseases). These cultivars were moderately to highly resistant to rusts and yield losses were mostly attributed to STB. Yield losses due to STB were calculated by subtracting the yield in the unsprayed plot from that in the sprayed plot. Based on these data, we predicted regional yield losses for each year with a linear model (fixed effects: year, region; random effects: wheat cultivar, trial). The national trend and the regional heterogeneity of yield losses, expressed in decitons per hectare, are shown in Fig. 1. This variable is denoted *P*_*it*_ hereafter, with *t* corresponding to the year and *i* to the region.

#### 2.1.4 Proportion of the total area under wheat farmed organically

The area under organically farmed wheat crops was recorded by AgenceBIO (the French national platform for the promotion and development of organic farming) and ARVALIS-Institut du Végétal. We collected regional data from 2007 onwards, and national data from 2004 onwards. The regional areas under organic wheat between 2004 and 2006 were assessed from the observed mean proportions of the regional area under organic wheat in subsequent years and from national data for 2004 to 2006. We used the regional proportion of wheat under organic farming in our models and calculated it by dividing the regional area under organic wheat by the total area under wheat in the same region, based on AGRESTE online data. The national trend and the regional heterogeneity of the area under organic wheat, expressed in hectares, are shown in Fig. 1. This variable is denoted *R*_*it*_ hereafter, with *t* corresponding to the year and *i* to the region.

### 2.2 Statistical modelling

We modelled the change in frequency for each resistance phenotype in French populations. The model took into account (*i*) the different phases of resistance dynamics (see below), (*ii*) the effects of previously described potential regional determinants and finally (*iii*) variability due to the sampling design (sampling date and wheat cultivar).

#### Phases in resistance dynamics

We distinguished three phases in resistance dynamics: “no resistance” (frequency equal to 0), “resistance selection” and “generalized resistance” (frequency equal to 100). During the “resistance selection” phase, observations were modelled with binomial random variables with a sample size of 100 (mean number of observed spores used to determine frequencies). The probabilities that a population was in the previous phases depended on the year *t*. These probabilities were referred as *π*_0*t*_, *π*_100*t*_ and (1 − *π*_0*t*_− *π*_100*t*_), respectively. Thus, *Y*_*itjkn*_, the *n*^*t*h^ frequency observed in region *i*, in year *t*, on cultivar *j* and at sampling date *k* followed a zero-and-one inflated binomial distribution (Equation 1).

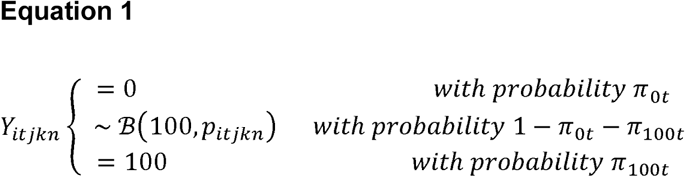

#### Resistance evolution

Using a logit transformation (Equation 2), the proportion *p*_*itjkn*_ of resistant phenotypes during resistance evolution was modelled by the regional dynamics *D*_*it*_ and the variability due to sampling design *ζ*_*itjkn*_.

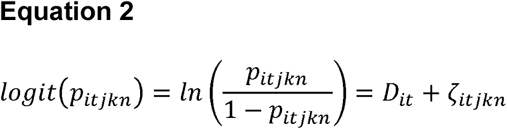

#### Regional dynamics

The change in resistance frequencies depended on regional-scale variables: fungicide use, yield losses and areas under organic farming. The regional dynamics *D*_*i*(*t*+1)_ in region *i* at year *t*+*1* was obtained by adding the regional dynamics of the previous year *D*_*it*_ to the additive effects of fungicide use *ϕ*_*it*_, yield loss *ρP*_*it*_ and wheat area under organic farming *κR_it_* in year *t* (Equation 3). *D*_*i*1_ is related to the logit of the initial resistance frequency in region *i*. We kept raw values for the fungicide use and area under organic wheat (*i*.*e*. neither centered nor reduced), while we centered values for the yield losses. Thus, the parameter β, the growth constant, corresponded to a continuous shift in resistance frequency in the absence of fungicides and refuges, with an average population size. This parameter will be interpreted as the relative apparent fitness.

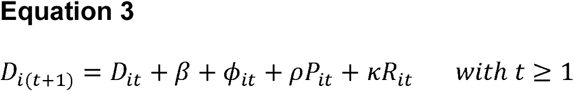

We derived two models from Eqn 3: one in which the use of fungicides is specified for each AI within MoAs, and another in which only the global use of each MoA (*i*.*e*. sum of AIs uses) is considered.

In the first model, **M**_***AI***_, the term *ϕ*_*it*_ from Equation 3 was defined as in Equation 4:

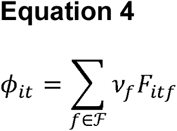

*F*_*itf*_ corresponds to the fungicide use for a specific AI *f*, in region *i* and year *t* (see section 2.1.2). The set *ℱ* included all AIs in a specific MoA associated with considered resistance: QoIs for the StrR phenotype, DMIs for the TriR phenotypes. We assumed that fungicide use positively selected resistant phenotypes over susceptible or less sensitive phenotypes. Thus, the parameters associated with fungicide use were, by definition, positive (*i*.*e. ν*_*f*_ ≥0), except for the TriR6 and TriR7-TriR8 phenotypes, which were not the most DMI-resistant phenotypes over their study period.

In the second model, **M**_***MoA***_, we considered: *F*_*it*._ = ∑ _*f*∈*F*_ *F*_*itf*_, the regional use of a given MoA. The term *ϕ*_*it*_ from Equation 3 was simplified and written: *ϕ*_*it*_ = *ν*\* *F*_*it*_. This model was run only for the StrR, TriMR and TriHR phenotypes, with, as above, the constraint *ν* ≥ 0.

#### Observation variability

The variability *ζ*_*itkln*_ of observations in Equation 2 was modelled with a mixed model (Equation 5).

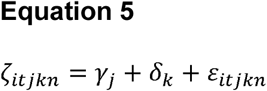

The parameter *γ*_*j*_ corresponds to the random effect of the 124 wheat cultivars (*γ*_*j*_ drawn from a centered Gaussian distribution with standard deviation *σ*_*y*_), considering an independent and constant effect of each cultivar over time. The parameter *δ*_*k*_ is the fixed effect of sampling date *k* (*i*.*e*. “S1” or “S2”, see section 2.1.1, with contrast *δ*_*S2*_ = 0). Finally, *ε*_*itjkn*_ is the overdispersion, modelled as a random individual effect with a mean of 0 and a standard deviation of *σ*.

### 2.3 Parameter expression

The explanatory variables had different units (*e*.*g*. proportion of wheat under organic farming *vs*. fungicide use). Moreover, the interpretation of the parameters of the zero-one-inflated logistic regression was not straightforward. We simplified the interpretation, by defining the *expected frequency difference* (EFD) for each variable and phenotype. The EFD described the frequency shift due to the mean value of this variable over a population with the mean resistance frequency. It is calculated as 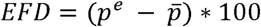 (*i*.*e*. the difference between *p*^*e*^, the *expected frequency*, and 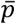, the *mean frequency* of the resistance phenotype in the data). The computation of the expected frequencies is detailed in Table 2.

**Table 2.**
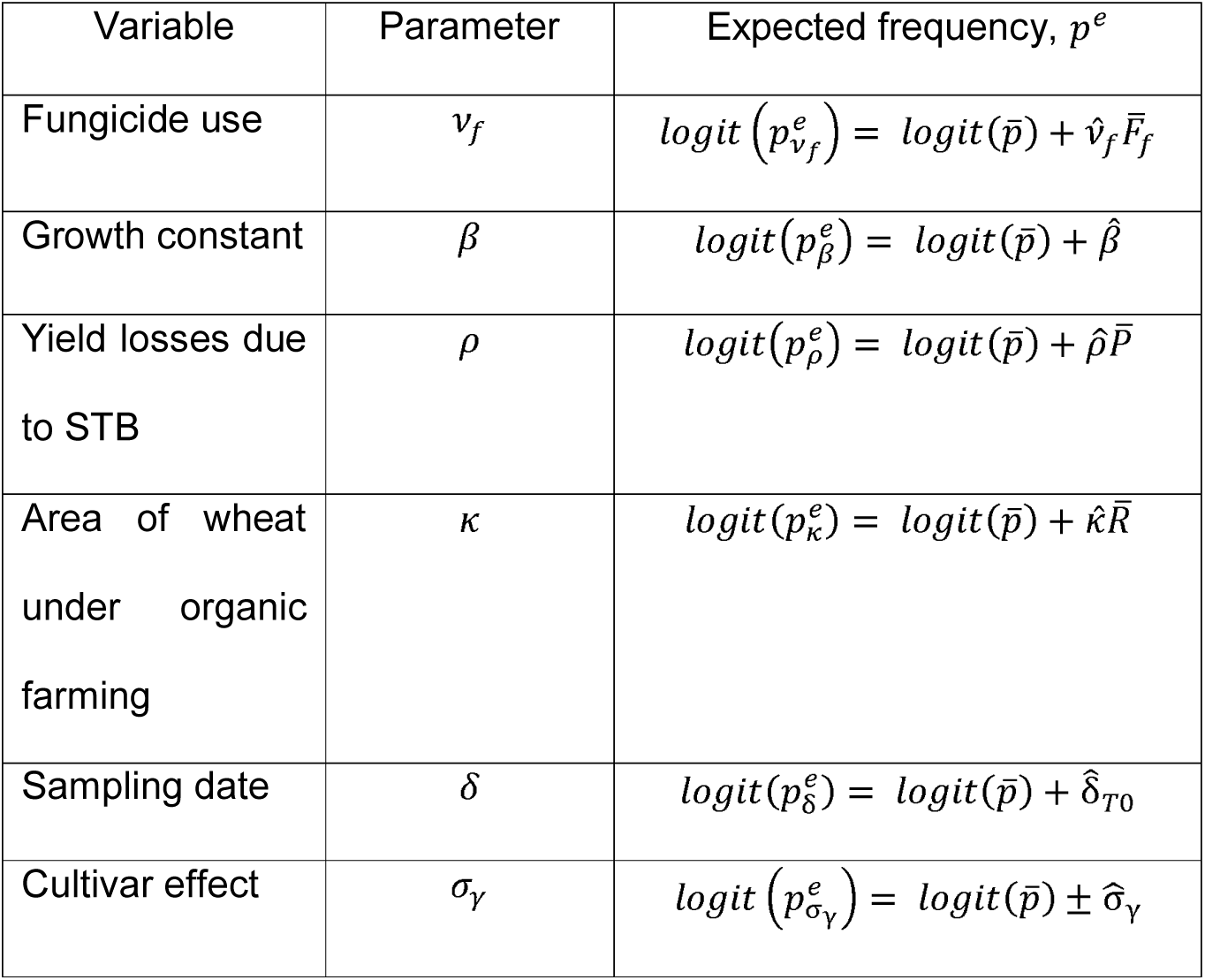
Computation of expected frequencies. Expected frequencies correspond to the resistance frequency obtained after exposing an average resistant population (*i*.*e*. at the frequency 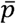 which is the mean frequency observed in data), to an average value of explanatory variables (*i*.*e*. 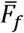 for the use of the fungicide *f*, 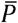 for yield losses induced by STB, 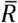 for organic wheat area). Each estimates are denoted with a hat 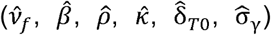 corresponds to the estimate extracted from models for the corresponding parameter. The difference between the expected and the mean frequencies isthe Expected Frequency Difference (EFD) for a given parameter.

| Variable | Parameter | Expected frequency, $p^e$ |
| --- | --- | --- |
| Fungicide use | $v_f$ | $\text{logit}(p_{v_f}^e) = \text{logit}(\bar{p}) + \hat{v}_f \bar{F}_f$ |
| Growth constant | $\beta$ | $\text{logit}(p_\beta^e) = \text{logit}(\bar{p}) + \hat{\beta}$ |
| Yield losses due to STB | $\rho$ | $\text{logit}(p_\rho^e) = \text{logit}(\bar{p}) + \hat{\rho} \bar{P}$ |
| Area of wheat under organic farming | $\kappa$ | $\text{logit}(p_\kappa^e) = \text{logit}(\bar{p}) + \hat{\kappa} \bar{R}$ |
| Sampling date | $\delta$ | $\text{logit}(p_\delta^e) = \text{logit}(\bar{p}) + \hat{\delta}_{T0}$ |
| Cultivar effect | $\sigma_\gamma$ | $\text{logit}(p_{\sigma_\gamma}^e) = \text{logit}(\bar{p}) \pm \hat{\sigma}_\gamma$ |

### 2.4 Bayesian analysis

Statistical analyses were performed with *R* software (R Development Core Team, 2008), in a Bayesian framework, with the *rjags* package (Plummer, 2013).

#### Prior and posterior densities

Non-informative prior distributions were used (Supporting Information, Eqn S1). Posterior distributions were estimated by Monte Carlo-Markov chain (MCMC) methods. Five MCMC chains were run, over 1 000 000 iterations, with a burn-in of 100 000 and a thinning every 1 000 for the variable selection phase (see the following section), followed by 500 000 iterations with a burn-in of 50 000 and a thinning every 500 for the final parameter estimation. Convergence was assessed with the Gelman and Rubin 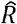 statistic (Gelman *et al*., 2004). Credible intervals of the highest posterior density were calculated from posterior densities with the *HDI* package (Dezeure *et al*., 2015). Parameter estimates were considered significant at the 5% level (or the 2.5% or 0.1% level), if their 95% credible interval (97.5% and 99.9%, respectively) did not contain the 0 value.

#### Variable selection

A selection procedure identified the relevant variables in each model for each resistance phenotype. We used a method based on indicator variables (Kuo & Mallick, 1998), in which each predictor was multiplied by a dummy variable with a prior distribution corresponding to a Bernoulli distribution with parameter *p* = 0.5. A predictor was retained in the model if the posterior expectation of its indicator variable was greater than 0.75 (thus, greater than its prior expectation of 0.5).

#### Variable weight

We assessed the influence of each explanatory variable *θ* by calculating its weight (*W*_*θ*_). The weight was *W*_*θ*_ defined as the ratio of *RSS*_*full*−*θ*_ to *RSS*_*full*_, where *RSS*_*full*_ and *RSS*_*full*−*θ*_ are the residual sums of squares of the full model (*i*.*e*. including all the explanatory variables selected by the variable selection procedure) and of this same model but without the explanatory variable *θ*, respectively. *RSS* should be minimal for the full model, so removing an explanatory variable should increase *RSS* : the more information *θ* contributes, the greater the increase in *RSS* and the higher the value of *W*_*θ*_. Conversely, if the information provided by *θ* is negligible, *RSS* is unaffected and *W*_*θ*_ is minimal (*i*.*e*. close to 1). In the result tables, we have calculated the relative weights by dividing individual variable weights by the sum of the weights of all variables.

#### Model comparison

For comparison of the M_*AI*_ and *M*_*MoA*_ models, we calculated the deviance information criterion, DIC (Plummer, 2013), and the coefficient of determination, *R*^2^ We also assessed the bias of the estimates using posterior predictive checks (Supporting Information, Eqn S2).

#### Predicted data

We computed predictions of the resistance frequencies for each phenotype for a given region *I*, and a given year *T* (Equation 6).

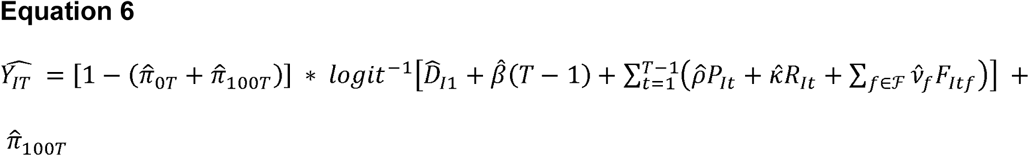

where parameter-hat are parameter estimates (*i*.*e*. their posterior mean). We therefore built maps of resistance status, for known initial frequencies, use of fungicides, yield losses and areas under organic wheat, from year 1 to year *T-1* in region *I*.

## 3 RESULTS

### 3.1 Overview of model fits

The convergence of the MCMC chain was satisfactory for all models (*i*.*e*. the Gelman and Rubin indicator 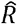 was below 1.1 for all parameters, in all models) and estimates were not biased (*PP*_*check*_ between 0.498 and 0.511).

After the selection procedure, no effect was retained for the following models: M_*AI*_ for the TriHR phenotype and M_*MoA*_ for the TriMR and the TriHR groups of phenotypes. The effects of fungicide use, yield losses, and areas of wheat under organic farming were not significant. As all estimates of the parameters of interest were equal to 0, we do not show and discuss the results of these models.

Finally, with Spearman’s method, a few significant correlations were found between some AI uses (*F*_*itf*_) in model inputs, but no significant correlation between estimates 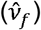 was found in model outputs (Supporting Information, Fig. S1). No correlation exceeded 0.4 in absolute value between AI uses and organic surfaces, nor between AI uses and yield losses.

### 3.2 Ranking of variable weight

Regional fungicide use appeared to be the major factor driving resistance evolution. In the M_*AI*_ models, which explicitly considered each AI, regional fungicide use systematically had the highest relative weight. For the StrR and TriMR phenotypes, it accounted for 87.4% and 72.6%, respectively. It accounted for 53.1% and 43.3% for the TriR6 and TriR7-TriR8 phenotypes, respectively (Table 3). For the M_*MoA*_ models, fungicide use, considered as the sum of AI uses within the same MoA, was also the major determinant of the StrR phenotype (associated with qualitative resistance to QoIs), accounting for 79.4% of the sum of variable weights (Table 4). For the TriMR and TriHR groups of phenotypes, no explanatory variables were selected for the M_*MoA*_ models.

**Table 3.**
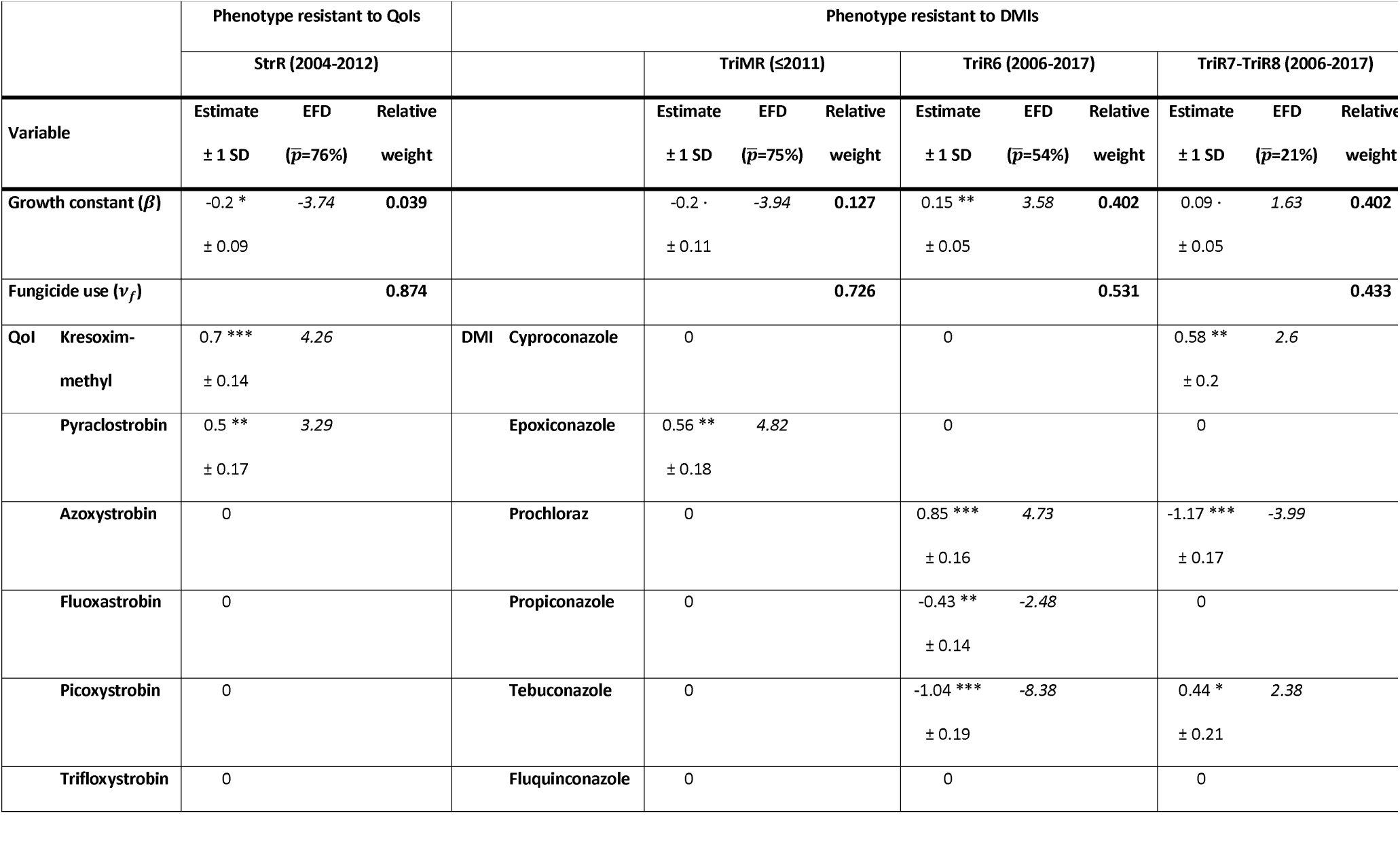
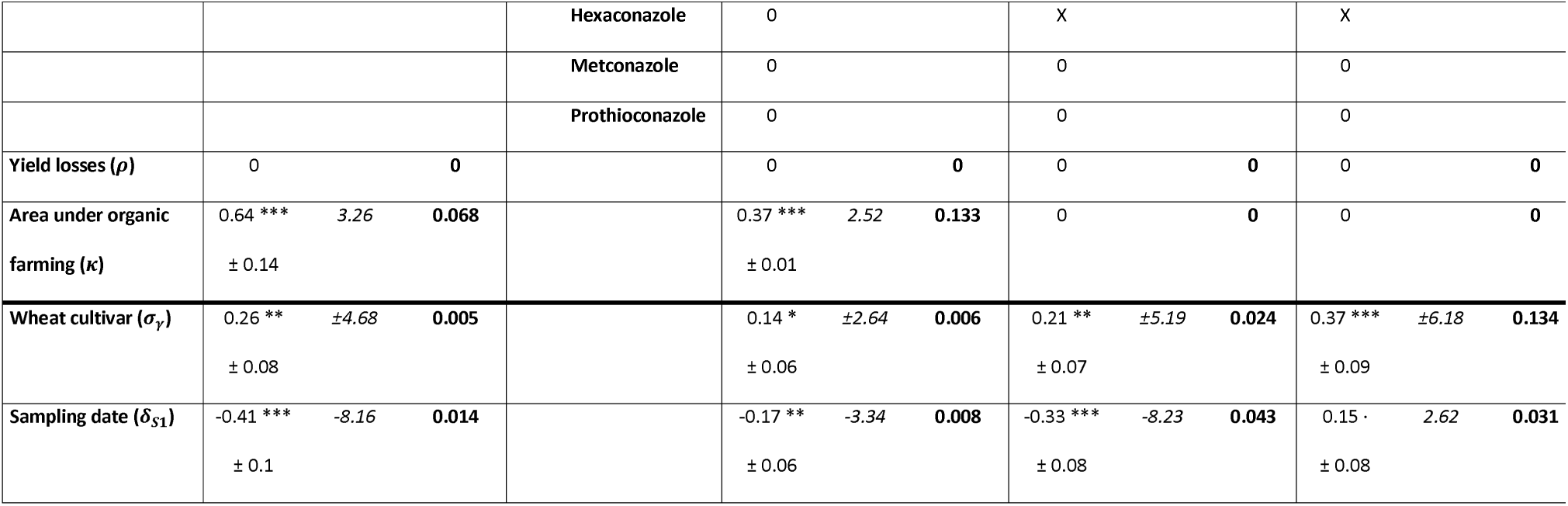
Estimates from the M_*AI*_ model for the StrR, TriMR, TriR6 and TriR7-TriR8 resistance phenotypes. For each phenotype, the three subcolumns show parameter estimates (*i*.*e*. posterior mean) and their variability (*i*.*e*. posterior standard deviation), expected frequency difference (EFD, *i*.*e*. the change in frequency due to the mean value of the variable on the reference population), and relative weights (*i*.*e*. the contribution of the variable to data variability). Note that the EFD therefore depends on the resistance frequency in the reference population 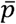, that we set at the average frequency observed in data. The significance thresholds are 0.1, 0.05, 0.025 and 0.001 denoted by “.”, “*”, “**” and “***”, respectively. Rows represent the different parameters of the models, ordered as follows: growth constant, regional scale explanatory variables (fungicide use, yield losses and area of wheat under organic farming) and local variation factors (wheat cultivar and sampling date). “0” means that the parameter was not selected during variable selection. “X” means that the variable was not considered in the model for the given phenotype. The results for the TriHR phenotypes are not shown as no explanatory variables were retained by the selection procedure.

**Table 4.** Estimates from the M_*MoA*_ model for the StrR phenotype. The three subcolumns show the parameter estimates (*i*.*e*. posterior mean) and their variability (*i*.*e*. posterior standard deviation), expected frequency difference (EFD, *i*.*e*. the change in frequency due to the mean value of the variable on the reference population), and relative weights (*i*.*e*. the contribution of the variable to data variability). Note that the EFD therefore depends on the resistance frequency in the reference population 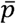, that we set at the average frequency observed in data. The significance thresholds are 0.1, 0.05, 0.025 and 0.001 denoted by “.”, “*”, “**” and “***”, respectively. Rows represent the different parameters of models, ordered as follows: growth constant, regional scale explanatory variables (fungicide use, yield losses and area of wheat under organic farming) and local variation factors (wheat cultivar and sampling date). “0” means that the parameter was not selected during variable selection. The results for TriMR and TriHR resistance phenotypes are not shown as no explanatory variable was retained by the selection procedure.

|  | StrR (2004-2012) |  |  |
| --- | --- | --- | --- |
| Variable | Estimate<br>$\pm 1$ SD | EFD<br>($\bar{p}=76\%$) | Relative<br>weight |
| Growth constant ($\beta$) | -0.17<br>$\pm 0.13$ | -3.23 | 0.041 |
| Fungicide use ($\nu$) | 1.07 ***<br>$\pm 0.26$ | 6.64 | 0.794 |
| Yield losses ($\rho$) | 0 | | 0 |
| Area under organic<br>farming ($\kappa$) | 0.78 ***<br>$\pm 0.16$ | 3.95 | 0.146 |
| Wheat cultivar ($\sigma_\gamma$) | 0.31 **<br>$\pm 0.09$ | $\pm 5.57$ | 0.009 |
| Sampling date ($\delta_{s1}$) | -0.35 ***<br>$\pm 0.1$ | -6.87 | 0.01 |

The growth constant was also an important parameter, albeit to a lesser extent. In M_*AI*_ models, the weight of the growth constant was lower than that of regional fungicide use by a factor of 5.7 times for TriMR phenotypes, 1.32 for the TriR6 phenotype, and 1.08 for the TriR7-TriR8 phenotypes (Table 3). For the StrR phenotype, the growth constant ranked third, with a weight lower than that of fungicide use by a factor of almost 20, for both models (Tables 3 and 4).

The proportion of the area under wheat farmed organically had a high relative weight for the StrR and TriMR phenotypes, but was not selected for TriR6 and TriR7-TriR8. For the StrR phenotype, its relative weight was ranked second, at 14.6% and 6.8% in the M_*MoA*_ and the M_*AI*_ models, respectively (Tables 3 and 4). For TriMR phenotypes, the relative weight of the wheat area under organic farming was about 13.3%, a value very similar to that for the growth constant (Table 3). Yield loss was systematically excluded during the selection procedure, for all models and phenotypes.

Sampling data and wheat cultivar, variables reflecting local variability, had only a low relative weight, with values always below 5%, except for the wheat cultivar variable for the TriR7-TriR8 phenotype, for which the value was 13.4% (Table 3).

### 3.3 Effect of variables at the regional scale

#### 3.3.1 Regional fungicide use

For the StrR phenotype, in the M_*MoA*_ model, the effect of the overall use of QoI fungicides was highly significant (*ν* = 1.07, *P* < 0.001) and the expected frequency difference (EFD) was estimated at 6.64%. Thus, a year of an average use of QoIs would have led to an increase of 6.64 frequency point on an average population composed of 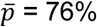 of StrR (Table 4).

In the M_*AI*_ model, two fungicides from the six QoI AIs were selected: kresoxim-methyl and pyraclostrobin. Their EFDs were similar: 4.26% (*ν* = 0.7, *P* < 0.001) and 3.29% (*ν* = 0.5, *P* < 0.025), respectively (Table 3). The M_*MoA*_ and M_*AI*_ models also had similar adequacies to data, according to their DIC (3480.2 and 3468.8 respectively) and *R*^2^ values (0.82 and 0.81, respectively).

For TriMR phenotypes, the effect of DMI use was estimated only in the M_*AI*_ model, as no explanatory variable was selected in the M_*MoA*_ model. One AI of the nine DMI fungicides was selected: epoxiconazole (*ν* = 0.56, *P* < 0.025) with an estimated positive EFD of 4.82% (with 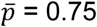, Table 3).

As the TriR6 and TriR7-TriR8 phenotypes were not the most resistant to DMI over the study period, the effect of fungicide use was not constrained to be null or positive, and was estimated only in the M_*AI*_ model. For the TriR6 phenotype, three AIs from the nine DMI fungicides were selected: prochloraz, with a positive EFD of 4.73% 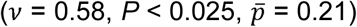, propiconazole with a negative EFD of -2.48% (*ν* = -0.43, *P* < 0.025) and tebuconazole with a negative EFD of -8.38% (*ν* = -1.04, *P* < 0.001). Thus, prochloraz use increased the frequency of TriR6, whereas tebuconazole counterselected TriR6 strains. For the TriR7-TriR8 phenotype, three AIs from the nine DMI fungicides were selected: cyproconazole and tebuconazole, with positive EFDs of 2.6% 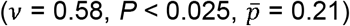 at 2.38% (*ν* = 0.44, *P* < 0.05), respectively, and prochloraz, with a negative EFD of -3.99% (*ν* = -1.17, *P* < 0.001). Prochloraz and tebuconazole clearly had opposite selection effects on TriR6 and TriR7-TriR8 phenotypes.

#### 3.3.2 Growth constant

The growth constant EFD quantified the change in resistance in the absence of regional fungicide use and unsprayed refuges, for a mean potential yield loss. It was therefore an extrapolation of the relative apparent fitness (referred to hereafter simply as fitness) of the phenotype considered (*i*.*e*. how much faster the phenotype would grew compared to the rest of the population (Hartl & Clark, 1997) in a year without fungicide treatment). A negative growth constant indicates a fitness cost, whereas a positive growth constant indicates a fitness gain. For the StrR phenotype, the growth constant provided an indication of the fitness of the resistant strains relative to the sensitive strains. The estimated fitness costs for this phenotype were similar in the M_*AI*_ (−3.74%; *β* = -0.2, *P* < 0.05; Table 3) and M_*MoA*_ (−3.23%; *β* = -0.17, P = 0.18; Table 4) models. DMIs selected a large diversity of phenotypes (TriLR, TriMR and TriHR groups), and no sensitive strains were detected during the study period. For TriMR phenotypes, the model was estimated with data from 2007 to 2011, when the frequency of the TriHR phenotype was still negligible (Fig. 1). Thus, the growth constant for TriMR phenotypes mostly compared their fitness with that of TriLR phenotypes. It was estimated at -3.94%, of borderline significance (*β* = -0.2, *P* < 0.1), and was associated with a relative weight of 12.7% (Table 3). The TriMR group included the TriR6 and TriR7-TriR8 phenotypes. For the TriR6 and TriR7-TriR8 strains, the model was estimated with data from 2006 to 2017. However, the TriHR phenotype has been non-negligible since 2014 (Figure 1). Thus, the growth constant for TriR6 strains compared their fitness with that of all the other phenotypes in the population: TriLR, TriR7-TriR8 and TriHR. The TriR6 growth constant was estimated at 3.58% (*β* = 0.15, *P* < 0.025; Table 3). This result may reflect a balance between a fitness cost of TriR6 relative to TriLR and TriR7-TriR8 strains, and a fitness gain relative to the TriHR phenotype. This rationale also applies to the apparent fitness gain of TriR7-TriR8, estimated at +1.63% (*β* = 0.09, *P* < 0.1; Table 3).

#### 3.3.3 Wheat area under organic farming

The proportion of the area under wheat management by organic farming methods increased resistance frequency, with an EFD estimated at 3.95% and 3.26% in the M_*MoA*_ and M_*AI*_ models, respectively, for the StrR phenotype (*κ* = 0.78 and 0.64, *P* < 0.001; Tables 3 and 4), and at 2.52% for the TriMR phenotype in the M_*AI*_ model (*κ* = 0.37, *P* < 0.001; Table 3). This variable was not selected for the TriR6 and TriR7-TriR8 phenotypes.

#### 3.3.4 Yield losses

The selection procedure did not retain the yield loss variable in any of the models.

### 3.4 Prediction maps

We mapped the predicted resistance frequencies for each year and for all phenotypes. Our maps predicting StrR phenotype dynamics were mostly based on regional QoI fungicide use and, to a lesser extent, the area under organic wheat, as explanatory variables (Fig. 2). Predictions mimicked the spatial propagation from the north to the south of France observed between 2004 and 2011 (as described by Garnault *et al*., 2019). Based on regional DMI use, our model accurately predicted (Fig. 3) the observed spatial partitioning of TriR6 strains, which were found mostly in the north east of France, and TriR7-TriR8 phenotypes, which were mostly localised in the south west (as described by Garnault *et al*., 2019).

**Fig. 2.**
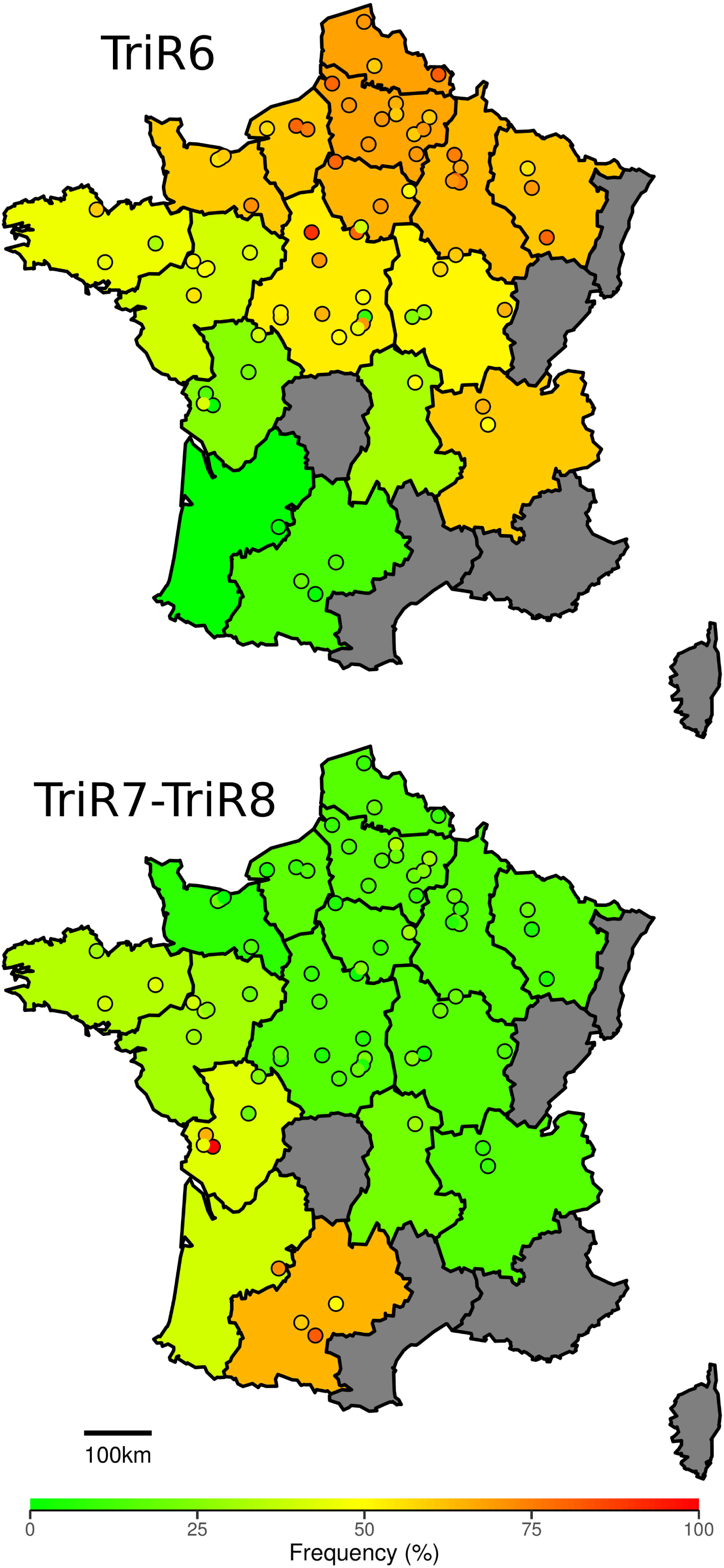
Maps of observed and predicted frequencies of the StrR resistance phenotype from 2004 to 2011. Real observations are represented by dots. The colour within the dot indicates the observed frequency in trials. The background map color shows the regional prediction from the M_*AI*_ model.

**Fig. 3.**
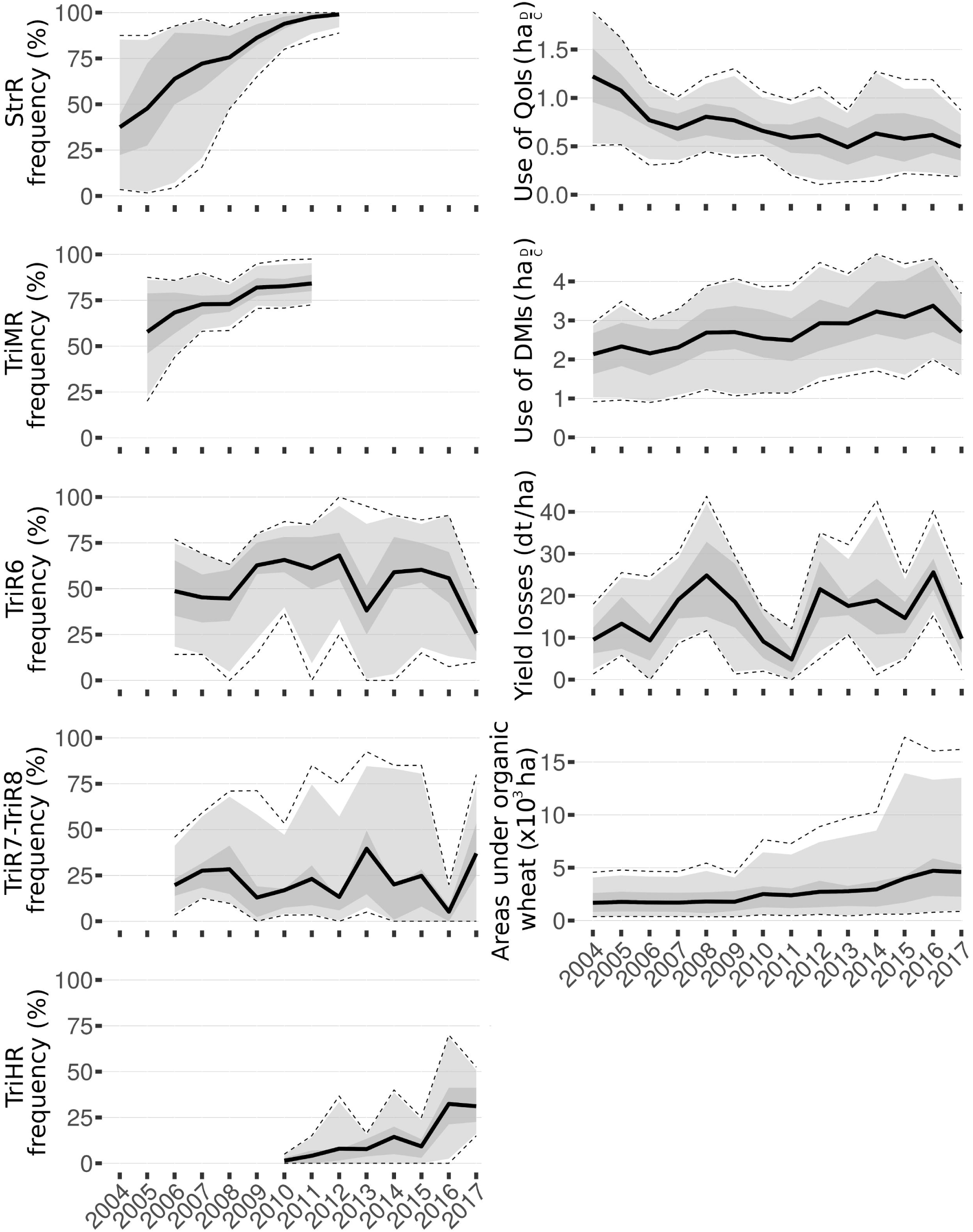
Maps of observed and predicted frequencies of TriR6 and TriR7-TriR8 resistance phenotypes in 2008. Real observations are represented by dots. The colour within the dot indicates the observed frequency in trials. The background map colour shows the regional prediction from the M_*AI*_ model. Top: TriR6. Bottom: TriR7-TriR8.

## 4 DISCUSSION

Here, we developed a model for identifying the determinants of fungicide resistance evolution at the regional scale. The candidate explanatory variables were regional fungicide use, pathogen population size (approximated by potential yield losses) and the fraction of refuges (approximated by the fraction of fields under organic wheat in a region). We analysed resistance frequencies in *Z. tritici* populations on winter wheat. Frequencies were monitored by the Performance trial network over the national territory of French (2004–2017; ~70 locations each year). We studied resistance against fungicides with two modes of action: QoIs, associated with qualitative resistance (StrR phenotype), and DMIs, associated with quantitative resistance (a continuum of multiple resistance phenotypes forming three main groups: TriLR, TriMR and TriHR, with low, medium and high levels of resistance to DMIs, respectively). The TriMR phenotypes encompassed two phenotypes: TriR6 and TriR7-TriR8.

### The regional use of fungicides is the main driver of resistance evolution at the plot scale

We demonstrated that regional fungicide use was a major determinant of the evolution of resistance. Fungicide use data at regional level, without taking into account the use of particular fungicides in particular fields, was sufficiently informative to explain resistance dynamics. Fungicide selection is, therefore, a large-scale process, and an understanding of the evolution of fungicide resistance requires consideration of the regional use of these compounds.

For qualitative resistance (e.g. StrR phenotype), the global use of a MoA (*i*.*e*. summed uses of the AIs of this MoA), can be considered as a good predictor. Distinguishing the effects of individual AIs within the MoA did not improve the fit of the model. By contrast, for quantitative resistance, it appeared to be necessary to consider each AI within the MoA separately, as they may select for different phenotypes (*e*.*g*. antagonist effects of prochloraz, tebuconazole on TriR phenotypes).

The AIs selected by the model were consistent with the history of fungicide use and/or with the patterns of cross-resistance described for some phenotypes. For instance, kresoxim-methyl was one of the first QoIs authorised in France (commercialised in 1997, www.ephy.anses.fr) and was selected to explain the evolution of StrR phenotypes, whereas more recent QoIs were not. The fungicides most used over the study period (*i*.*e*. epoxiconazole for DMIs, pyraclostrobin and kresoxim-methyl for QoIs; Supporting Information, Fig. S2-S3), were also selected in models. The large-scale effect of AIs was also linked to the cross-resistance pattern of the phenotypes. The estimated selection effect of epoxiconazole was consistent with resistance factors (RFs) of TriMR phenotypes to epoxiconazole, being higher than those of TriLR phenotypes which were the second most frequent resistance phenotype during this period (RFs described in Supporting Information, Table S1 from Leroux & Walker, 2011). The selection for TriR6 and counterselection for TriR7-TriR8 phenotypes induced by the prochloraz were consistent with RFs (6.7 and <1.5, respectively), as well as the effect of counterselection for TriR6 and selection for TriR7-TriR8 phenotypes by tebuconazole (RF=74 and 91, respectively for TriR6 and TriR8, TriR7 being unfrequent (Huf *et al*., 2018)). Propiconazole and cyproconazole were often used together as a mixture, so were strongly correlated (Supporting Information, Fig. S1). The effects of counterselection for TriR6 phenotypes by propiconazole and of selection for TriR7-TriR8 by cyproconazole may indicate that their mixture globally promoted the selection of TriR7-TriR8 strains over TriR6 strains. Again, this is consistent with RFs to propiconazole (35 and 54, respectively for TriR6 and TriR8 strains) and to cyproconazole (11 and 13, respectively). These findings also confirm that the laboratory characterisation of strains can be good predictor of resistance evolution in the field, if properly used, as reported by Blake *et al*. (2018).

Our findings highlight the importance of defining precise resistance profile phenotypes. Indeed, the model fitted slightly better the frequencies on more homogeneous phenotypes (R^2^ = 0.48 for TriMR vs. 0.56 for TriR6 and TriR7-TriR8 phenotypes). In addition, no significant effect of the different DMIs was found for the TriHR phenotype group. Indeed, TriHR strains encompass multiple heterogeneous phenotypes, resulting from combinations of target alteration, target overexpression and enhanced efflux, as resistance mechanisms (Leroux & Walker, 2011; Huf *et al*., 2018). The tremendous diversity and redundancy of phenotypes observed in the field, especially for TriHR phenotypes, made it possible to classify strains only approximately. There is therefore a need to develop molecular tools for quantifying genotypes rather than phenotypes. Multi-trait high-throughput genotyping should quantify resistance frequency more accurately and ultimately lead to improvements in our ability to predict resistance evolution.

### The proportion of the wheat area under organic farming may be still too limited to mitigate the evolution of resistance via a refuge effect

Wheat areas managed under organic farming systems are not treated with synthetic fungicides. As some conventional fields may be unsprayed with a given MoA, organic areas represent a lower bound of the surfaces that were unsprayed with this MoA (only true for QoIs and SDHIs, since DMIs use is near universal in conventional wheat farming).They could act as refuges for “wild” susceptible or less resistant individuals, which may reproduce, delaying the evolution of resistance by a dilution effect in mobile species (Gould, 2000). But refuges may also provide a heterogeneous environment, promoting sink-source dynamics: selection-free wheat areas acting as a source of susceptible strains that migrate towards areas farmed conventionally. These opposite effects have been described in studies of the resistance to transgenic crops expressing *Bacillus thuringiensis* (*Bt*) toxins, where non-*Bt* crops may act as refuges diluting the selection (Huang *et al*., 2011), or promoting the evolution of resistance (Caprio, 2001).

The refuge effect for *Z. tritici* is, theoretically, weak due to the haploid nature of this organism (Shaw, 2009), but this has never been studied experimentally. We did not validate the beneficial effect of refuges, approximated by the area under wheat farmed organically. On the contrary, we estimated that the selection of StrR and TriMR phenotypes would increase with wheat areas under organic farming. However, the weight of this explanatory variable remained much lower than that for regional fungicide use. In addition, the evolution of surfaces under organic wheat was very different between South and North of France, with both higher proportions and faster increase in Southern regions (Supporting Information, Fig. S4). Therefore, the effect of refuges remains difficult to distinguish from other potential factors reflecting differences between Northern and Southern agriculture (e.g. drier climate, higher proportion of durum wheat, in Southern regions). This effect was not found significant for the other phenotypes.

According to Huang *et al*. (2011), three conditions must be satisfied for refuge strategies to be successful: selection at “high dose”, a very low initial frequency of resistance, and sufficient refuge areas located nearby. In our study, initial frequencies of resistance (*i*.*e*. at the very beginning of our study) were already quite high (up to 85% in some regions, Fig. 1). In addition, the area under organic farming may still be too small within the landscape (generally less than 1% until 2010). Nevertheless, its proportion has steadily increased since the 2010s (Fig. 1). The effect of refuges may become detectable in those areas, and should be investigated further, particularly for emerging resistance phenotypes.

### The growth constant reveals a relative fitness penalty of phenotypes

The growth constant represents the evolution of phenotypes in the absence of fungicide treatment. It represents the apparent fitness of the phenotype relative to that of the susceptible phenotype (or other resistant phenotypes, in the case of quantitative resistance). The term “apparent” is used because this quantification takes place in current crop conditions. The fitness cost of resistance to drugs is known to be a key parameter driving effective anti-resistance strategies (Andersson & Hughes, 2010; Melnyk *et al*., 2015; Mikaberidze & McDonald, 2015). If there is no fitness cost, regardless of the strategy used, resistance will always be selected irreversibly. However, it remains difficult to infer the global fitness cost of a mutation throughout the entire life cycle of a pathogen (Hollomon, 2015).

We inferred an apparent relative fitness penalty for StrR phenotypes, resulting in an annual decrease of 3.74%. This was consistent with the fitness cost described by Hagerty & Mundt (2016) using virulence comparison tests. The impact of cyp51 alterations, leading to TriR phenotypes, on fitness is often invoked to explain the evolution of azole resistance in *Z. tritici* populations (Cools *et al*., 2013; Blake *et al*., 2018) but it has not been quantified as yet. Our model is consistent with these assumptions as it also inferred an apparent fitness penalty for the TriMR group (−3.94% per year). Decreases in the use of QoI and DMI fungicides, and/or the implementation of strategies favouring the expression of a resistance cost, may help to slow resistance evolution. From our estimates, we can extrapolate a theoretical equilibrium between resistance cost and selection, which could have led to a null growth of the StrR and TriMR phenotypes by reducing by 50% the use of QoIs and 18% for DMIs. Despite the resistance cost inferred for StrR phenotypes, their frequency remains high in French populations (> 95%), most probably due to QoIs residual use against rusts and *Fusarium* head blight (Supporting Information, Fig. S2).

For the TriR6 and TriR7-TriR8 phenotypes, the growth constant is an integrative value, as these phenotypes were studied alongside more susceptible phenotypes (TriLR before 2010) and more resistant phenotypes (TriHR after 2010). The positive growth constants we estimated for these two phenotypes may indicate a fitness benefit of TriMR relative to TriLR phenotypes and/or fitness cost of TriHR (*i*.*e*. also fitness benefit of TriMR relative to TriHR phenotypes). Our model could be extended to determine the relative fitnesses of each phenotype in cases of quantitative resistance. The global informative indicator provided by our model could be used to guide the design of optimal large-scale fungicide deployment strategies.

### The yield losses caused by STB do not affect resistance evolution

Population size, a major parameter in population adaptation (Good *et al*., 2012), is generally positively correlated with resistance evolution (Weber & Diggins, 1990; Anderson, 2005; zur Wiesch *et al*., 2011). Indeed, a large population size increases the number of mutants generated and decreases genetic drift (Linde *et al*., 2002). We inferred this effect using potential yield losses caused by STB as a proxy, but no significant effect was detected for any resistance phenotype. Population size may not be limiting for resistance evolution in *Z. tritici*, since the number of individuals remains important even in years with little disease (Zhan *et al*., 2001; Mikaberidze *et al*., 2017), particularly when considering large scales. Population size may not be described accurately enough as our proxy variable also depends on the timing of infection (Shaw & Royle, 1993) and on stubble management (McDonald & Mundt, 2016).

### Predicting resistance evolution over years

Prediction maps can be computed from our model, using only the initial regional frequencies of the phenotypes, the history of fungicide use (between the initial year and the year to be predicted) and the history of area under organic farming. Predictions for the StrR phenotype from 2004 to 2011 highlighted the same colonisation front structure from the North to the South of France (Fig. 2) as reported by Garnault *et al*. (2019) albeit the regions are assumed to be fully independent. Besides, the way StrR propagated from North to South may indicate that integrating regional interdependencies in the model would improve its performance. However, this would increase its complexity and require more data to estimate inter-regional fluxes. Predictions for the TriR6 and TriR7-TriR8 phenotypes also yielded stable spatial distributions between North-East and South-West France (Fig. 3), as previously observed in Garnault *et al*. (2019).

Further analysis will be required to assess the prediction quality of the model. Nevertheless, this finding supports the global validity of our model and paves the way for an original approach to predicting resistance evolution in a heterogeneous landscape.

### Conclusion

We developed a model of resistance dynamics, which identified regional fungicide use as the major determinant of fungicide resistance evolution, and the area under organic farming as a much weaker explanatory variable. We estimated the apparent relative fitness of resistant phenotypes, a key parameter for the development of sustainable resistance management strategies. We also identified AIs whose use drove resistance evolution.

As variability of resistance evolution is mainly explained by the regional intensity of selection induced by fungicides (without information on local application strategies), concerted collective action on the regional or national fungicide use could improve control of the evolution of resistance, especially in the most at-risk regions.

## 5 ACKNOWLEDGEMENTS

We acknowledge the two anonymous reviewers for their helpful comments. This work was funded by the INRAE SMaCH metaprogramme (FONDU project) and by Bayer Crop Science (PhD scholarship of M. Garnault; CIFRE no. 2016-0695). We warmly thank all partners of the Performance network for providing samples since 2004, and Arvalis-Institut du Végétal for supervising the network. We are grateful to the INRAE MIGALE bioinformatics facility (MIGALE, INRAE, 2020. Migale bioinformatics Facility, doi: 10.15454/1.5572390655343293E12) for providing computing resources. We thank Eva Lacarce (AgenceBio) and Phillipe Viaux (Arvalis-Institut du Végétal) for providing information about the historical changes in the area under organic wheat, and Clarisse Payet (Bayer Crop Science) for providing data on fungicide use in France and for helpful discussions. BioGER benefits from the support of Saclay Plant Sciences-SPS (ANR-17-EUR-0007).

## 6 AUTHOR CONTRIBUTIONS

GC was responsible for Performance network administration and supervision and for establishing the dataset. PL, ASW and CD carried out laboratory manipulations on the network samples to determine resistance frequencies in *Z. tritici* populations. FC and OD were involved in the development of the methodology and provided theoretical support for statistical aspects for data analyses. MG handled data, coded the model, performed statistical analyses and generated most of the results (figures, maps and tables) during his PhD work. The article was written by MG, FC, ASW and OD.

