## Supplemental Information 1 for "Large-scale study validates that regional fungicide applications are major determinants of resistance evolution in the wheat pathogen *Zymoseptoria tritici* in France"

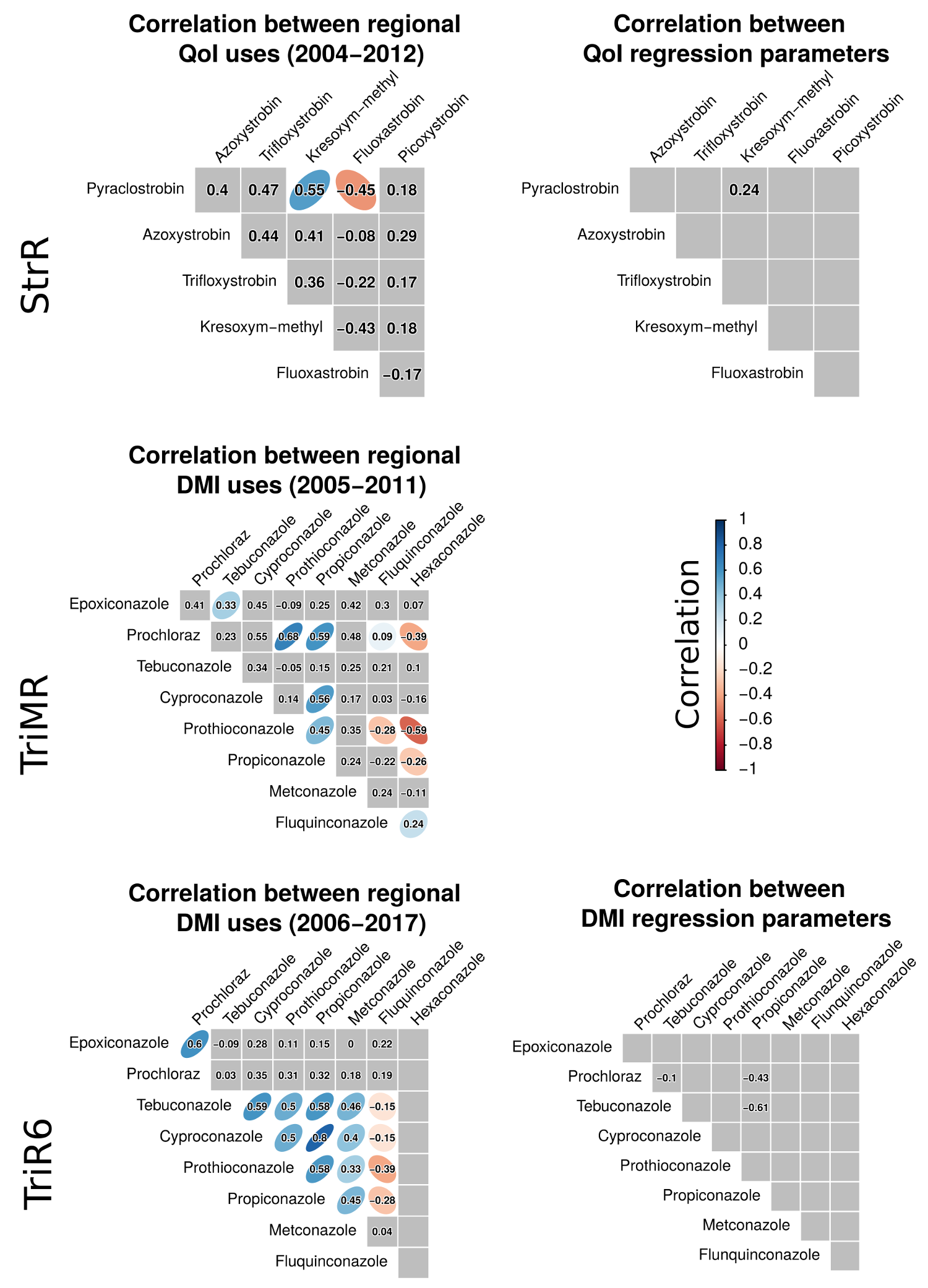

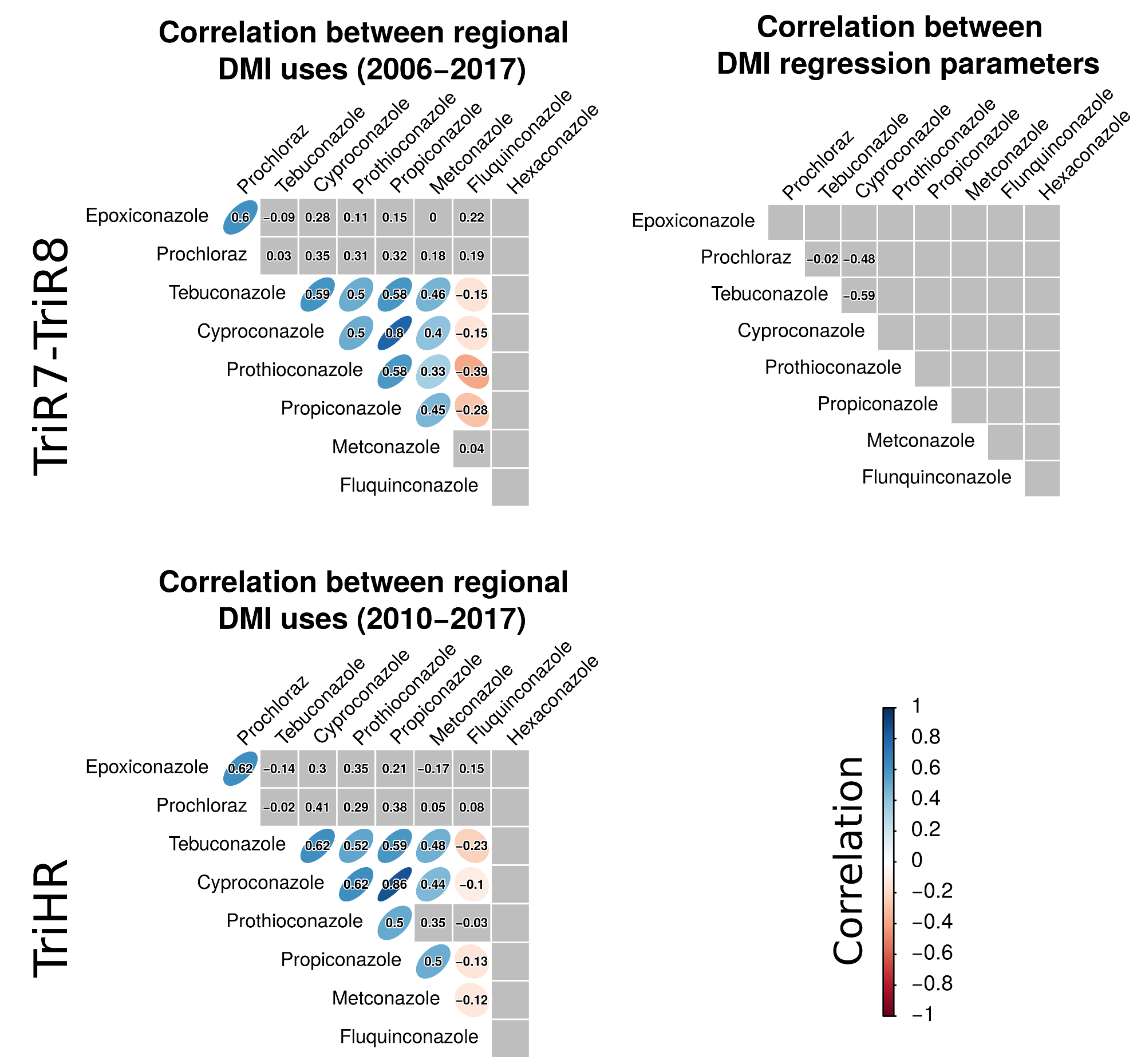

**Fig. S1** Correlation between the use of AIs at the regional scale (left), and between the posterior distributions of the associated regression parameters in M*_AI_* models (right).

Correlations were calculated with the Spearman method. A blue ellipse indicates a positive correlation, whereas a red ellipse indicates a negative correlation, a grey background indicates that the correlation was not significant at the 5% confidence level. Numbers correspond to the intensity of the correlation: the closer the absolute value of these values is to 1, the stronger the correlation. From top to bottom: AI use and their posterior distributions for the StrR, TriMR, TriR6, TriR7-TriR8 and TriHR phenotypes. No correlation between posterior distributions is shown for TriMR as only one AI was selected by the model. No correlation between posterior distributions is shown for TriHR strains as no AI was selected by the model.

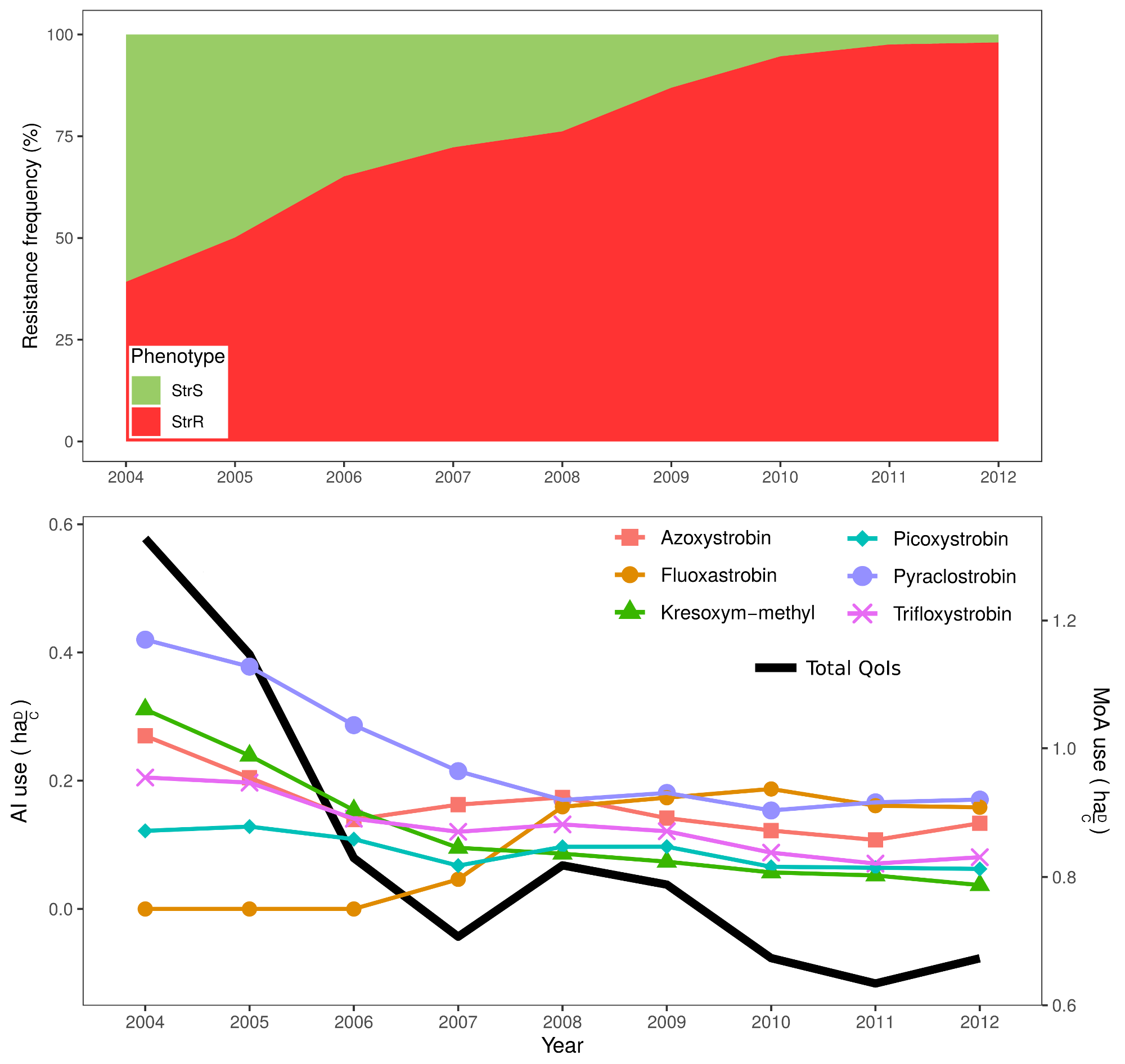

**Fig. S2** Patterns of change in national AI use and resistance in populations: QoIs / StrR.

Top: Change in the frequency of the StrR phenotype (specific resistance to QoIs) and the corresponding sensitive phenotype, StrS, in French *Z. tritici* populations between 2004 and 2012. Bottom: Change in QoI use (expressed in ${ha}_{\frac{D}{C}}$, *i.e.* mean number of times each active substance was used in sprays over a cropping season, regardless of the dose used) for azoxystrobin, fluoxastrobin, kresoxym-methyl, picoxystrobin, pyraclostrobin, trifloxystrobin and for the global QoI mode of action (sum of AI uses).

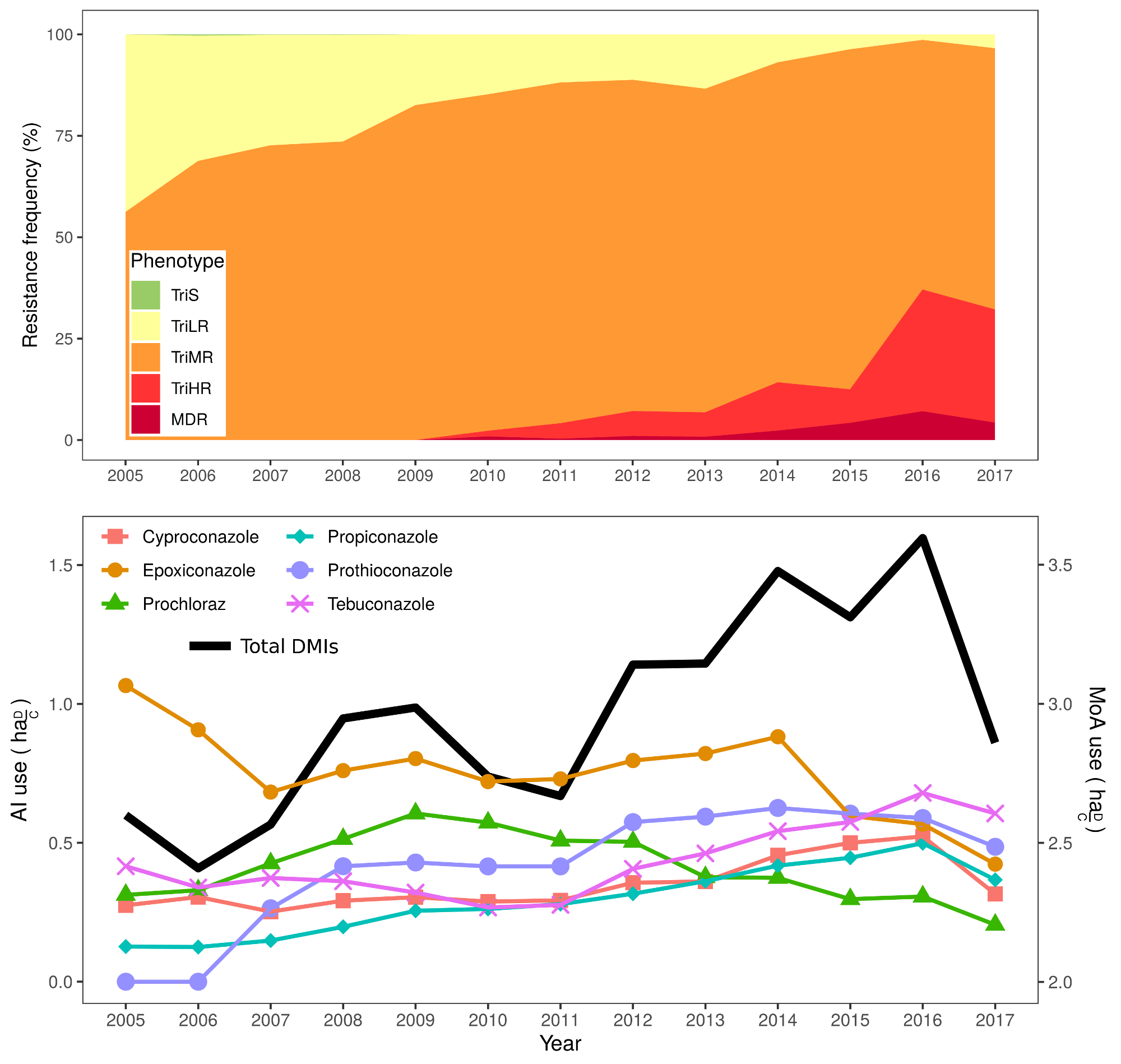

**Fig. S3** Changes in national AI use and resistance in populations: DMIs / TriRx.

Top: Change in the frequency of the TriR phenotypes (specific resistance to DMIs encompassing, TriLR, TriMR and TriHR strains), the corresponding sensitive phenotype, TriS, and the MDR phenotype, in French *Z. tritici* populations between 2005 and 2017. Bottom: Change in DMI use (expressed in ${ha}_{\frac{D}{C}}$, *i.e.* mean number of times each active substance was used in sprays over a cropping season, regardless of the dose used) for cyproconazole, epoxiconazole, prochloraz, propiconazole, prothioconazole, tebuconazole and for the global DMI mode of action (sum of AI uses).

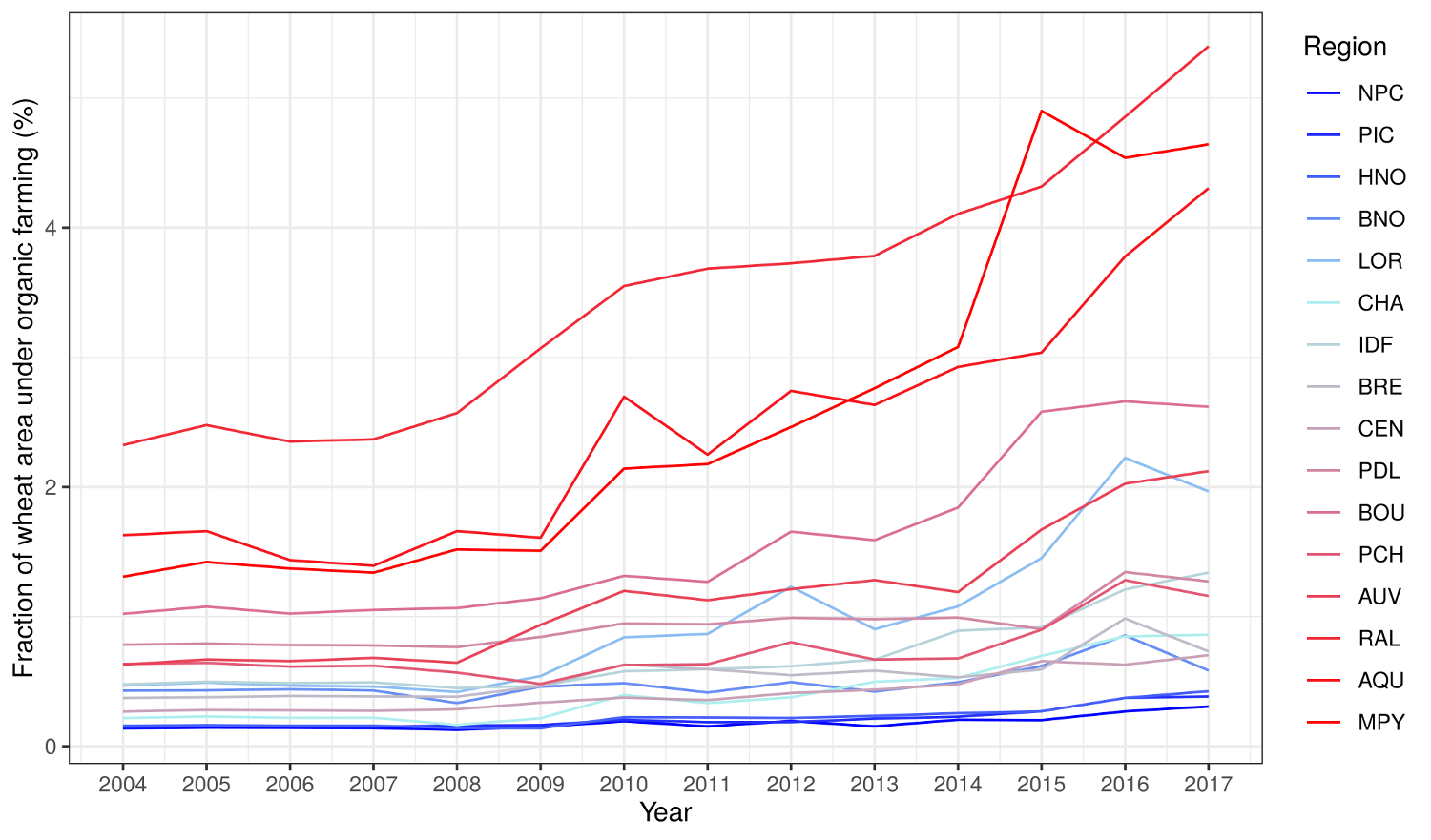

**Fig. S4** Temporal evolution of the fraction of surfaces under organic wheat in French regions.

The fraction is computed relatively to the total surfaces cropped with wheat. In the legend, regions are organised on a north (blue) to south (red) gradient. Region acronyms are: ALS, Alsace; AUV, Auvergne; AQU, Aquitaine; BNO, Basse-Normandie; BOU, Bourgogne; BRE, Bretagne; CEN, Centre, CHA, Champagne-Ardennes; FCO, Franche-Comté; HNO, Haute-Normandie; IDF, Ile-de-France; LAR, Languedoc-Roussillon; LIM, Limousin; LOR, Lorraine; MPY, Midi-Pyrénées; NPC, Nord-Pas-de-Calais; PCH, Poitou-Charentes; PDL, Pays de la Loire; PIC, Picardie; RAL, Rhône-Alpes.

**Eqn S1** JAGS script used to run M*_AI_* and M*_MoA_* models.

*model{*

*############*

*# Likelihood #*

*############*

*for(i in 1:n){*

*y[i] ~ dbin(P[i], 100)*

*P[i] <- p0[i] * 0 + pp[i] * p[i] + p100[i] * 1*

*p0[i]	<- equals(inflated[i], 1)*

*pp[i]	<- equals(inflated[i], 2)*

*p100[i] <- equals(inflated[i], 3)*

*inflated[i] ~ dcat(pi[year[i], ])*

*p[i] <- exp(eta[i])/(1+exp(eta[i]))*

*eta[i] <- mu[i] + epsilon[i]*

*mu[i] <- National.F0 + Regional.F0[region[i]] +*

*sum(Selection.Pressure[i, 1:n_fungicide]) +*

*Cte.Growth * time[i] +*

*Pop.Size * pop.size[i] +*

*Refuges * refuges[i] +*

*Sampling[sampling[i]] +*

*abs(Sigma.Cultivar) * Cultivar[cultivar[i]]*

*epsilon[i] ~ dnorm(0, Over.Dispersion)*

*for(j in 1:n_fungicide){*

*Selection.Pressure[i, j] <- Fungicide[j] * fungicide[i, j]*

*}*

*}*

*########*

*# Priors #*

*########*

*for(i in 1:3){*

*alpha[i] <- 1/2*

*}*

*for(i in 1:n_year){*

*pi[i, 1:3] ~ ddirch(alpha[])*

*}*

*Sigma.Over.Dispersion ~ dnorm(0, 1)T(0,)*

*Over.Dispersion <- 1/(Sigma.Over.Dispersion^2)*

*National.F0 ~ dnorm(0, 1.0E-1)*

*Sigma.Regional.F0 ~ dnorm(0, 1)T(0,)*

*InvVar.Regional.F0 <- 1/(Sigma.Regional.F0^2)*

*for(i in 1:n_region){*

*Regional.F0[i] ~ dnorm(0, InvVar.Regional.F0)*

*}*

*Cte.Growth ~ dnorm(0, 1.0E-1)*

*for(i in 1:n_fungicide){*

*Fungicide[i] ~ dnorm(0, 1.0E-1)*

*}*

*Pop.Size ~ dnorm(0, 1.0E-1)*

*Refuges ~ dnorm(0, 1.0E-1)*

*Sampling[1] ~ dnorm(0, 1.0E-1)*

*Sampling[2] <- 0*

*Sigma.Cultivar ~ dnorm(0, 1)*

*for(i in 1:n_variete){*

*Cultivar[i] ~ dnorm(0, 1)*

*}*

*}*

**Equation S2** Estimation bias indicator: ${PP}_{check}$

$${PP}_{itjkn}^{check}=P(y_{itjkn}-y_{itjkn}^{rep}<0 | Y)$$

We assessed the bias of the estimation using posterior predictive checks (Gelman *et al.*, 2004). Replicated data ($y_{itjkn}^{rep}$) generated during the MCMC algorithm from the model posterior densities were compared to observed data ($y_{itjkn}$). The mean value of ${PP}_{itjkn}^{check}$, where $Y$ is the vector of observations, is denoted ${PP}_{check}$. A non-bias estimation is achieved when ${PP}_{check}=0.5$, *i.e.* with equal probabilities of over- and under-estimation.

**Table S1** Resistance factors (RFs) for DMIs for the different TriR phenotypes, estimated from the germ tube elongation of *Z. tritici* field isolates on solid medium

|  | **TriLR** | | **TriMR** | | | **TriHR** | |
| --- | --- | --- | --- | --- | --- | --- | --- |
| **Fungicide** | **TriR4** | **TriR5** | **TriR6** | **TriR7** | **TriR8** | **TriR9** | **TriR11** |
| Prochloraz | 6.7 | 15 | **6.7** | **1.5** | **0.8** | 66.7 | 22.2 |
| Cyproconazole | 4.3 | 8.5 | 11.2 | **7.6** | **13.1** | 16.3 | 34.7 |
| Epoxiconazole | 5 | 8.5 | ***25.5*** | ***11*** | ***23*** | 29.9 | 60 |
| Fluquinconazole | 6.5 | 14.2 | 20.3 | 14.5 | 22.6 | 75.8 | 103.2 |
| Hexaconazole | 5.6 | 4.4 | 8.9 | 6.7 | 8.9 | 8.9 | 11.1 |
| Metconazole | 10 | 8 | 15.5 | 10 | 17.5 | 23.6 | 30 |
| Propiconazole | 12.2 | 20.5 | **35.1** | 27 | 54.1 | 70.9 | 54.1 |
| Tebuconazole | 18.2 | 1.8 | **74.5** | **51.8** | **90.9** | 12 | 18.2 |
| Prothioconazole^a^ | 3.8 | 5.3 | 7.8 | 7 | 7.3 | 16.3 | 25 |

Mean profiles were calculated from the data provided in Leroux & Walker, 2011. Only AIs used in models are shown. For each phenotype, bold values correspond to AIs selected in the M*_AI_* model. The values for epoxiconazole are shown in italics because this AI was not selected for either TriR6 or TriR7-TriR8 phenotypes separately, but was selected for the TriMR phenotype group (encompassing the TriR6 and TriR7-TriR8 phenotypes). For the TriLR phenotype group, we show RFs only for TriR4 and TriR5 strains, as these were the most common in France (Stammler *et al.*, 2008; Leroux & Walker, 2011; Stammler & Semar, 2011). Similarly, for the TriHR phenotype group, we show RFs only for the TriR9 and TriR11 strains, which were the most common in France (Leroux & Walker, 2011; Huf *et al.*, 2018).

^a^RFs measured for prothioconazole-desthio, the active metabolite of prothioconazole, may be greater than those for prothioconazole.

**Supporting Information References:**

**Gelman A, Carlin JB, Stern HS, Rubin DB**. **2004**. Bayesian data analysis 2nd edn Chapman & Hall. *CRC, Boca Raton FL*.

**Huf A, Rehfus A, Lorenz KH, Bryson R, Voegele RT, Stammler G**. **2018**. Proposal for a new nomenclature for CYP 51 haplotypes in Zymoseptoria tritici and analysis of their distribution in Europe. *Plant Pathology* **67**: 1706–1712.

**Leroux P, Walker A-S**. **2011**. Multiple mechanisms account for resistance to sterol 14α-demethylation inhibitors in field isolates of Mycosphaerella graminicola. *Pest management science* **67**: 44–59.

**Stammler G, Carstensen M, Koch A, Semar M, Strobel D, Schlehuber S**. **2008**. Frequency of different CYP51-haplotypes of Mycosphaerella graminicola and their impact on epoxiconazole-sensitivity and-field efficacy. *Crop Protection* **27**: 1448–1456.

**Stammler G, Semar M**. **2011**. Sensitivity of Mycosphaerella graminicola (anamorph: Septoria tritici) to DMI fungicides across Europe and impact on field performance. *EPPO Bulletin* **41**: 149–155.
